# An efficient method for visualizing the plaques of *Bdellovibrio bacteriovorus*

**DOI:** 10.1101/2024.06.24.600463

**Authors:** Qian Zhao, Jiangong Xu, Kui Zhu

## Abstract

*Bdellovibrio bacteriovorus* preys upon other Gram-negative bacteria. This obligate predator is recognized as a living antibiotic to control the rising problem of antibiotic resistance. Plaque forming units (PFU) counting is commonly used to determine the viable numbers of *B. bacteriovorus*. However, nearly 3-days incubation is always necessary for getting the single, obvious plaques on the double layer agar plate. This time-consuming procedure greatly impedes the purification and enumeration efficiency of *B. bacteriovorus*. In this work, we evaluated the advantages of fluorescent prey on the plaque visualization of the predator. Our study reveals that plaques of the *B. bacteriovorus* have already formed before they could be monitored by eyes on the double layer agar plate. The regular single plaques are easily observed under the enhanced background of fluorescent prey lawn in 1.5 days, reducing nearly half of the time consumption in the purification and enumeration of *B. bacteriovorus*. In addition, it also provides some paradigms for modeling and testing the subtle predation process on the solid medium.

**IMPORTANCE:** The viability of predator *Bdellovibrio bacteriovorus* is typically suggested by the lysis of the prey bacteria on the double layer agar plate. However, long period incubation is necessary to get single obvious plaques of *B. bacteriovorus*. Here, we found that plaques are already generated before it can be monitored by eyes. The plaques are obvious on the enhanced background of the double layer agar plate in the presence of fluorescent prey under UV light. We confirmed that the utilization of fluorescence labeled prey bacteria reduces the time spent on the isolation, purification, and enumeration of the predator *B. bacteriovorus* on the double layer agar plate.

## INTRODUCTION

*Bdellovibrio bacteriovorus* was firstly isolated from soil by *Heinz Stolp* in 1962. This fascinating predator can collide, invade, and kill diverse Gram-negative bacteria, such as *Salmonella, Enterobacteriaceae*, and *Pseudomonas*^1, 2^. The potential application as “living antibiotic” has gained huge attention in response to the increasing concern of antibiotic resistance^3-5^. Due to the small size of *Bdellovibrio* (0.2-0.5 by 0.5-2.5 μm), the bacterial density cannot be measured through the cultural turbidity reliably. Predation, or the viability of predator is typically indirectly reflected by the reduction in cell density or by gradient dilution and viability counts of the prey bacteria. With the progress of new genetic tools and techniques, luminescent and fluorescent prey are applied to indicate changes in the predator viability and prey population^6-9^. Other methods such as real-time quantitative PCR^10, 11^, flow cytometry^12^ and fluorescence visualization^13, 14^ are utilized to detect the presence and viability of the predator. Even these rapid methods are described, PFU (plaques forming unit) counting on the double-layer agar plate is still the conventional, economic, and standard methods for detect and quantify the predator viability and biomass^13, 15, 16^. As plaques are generally visible at 3 days by eyes on the double layer agar plate, the long growth period greatly hampered its application on the enumeration of predatory populations. Besides, to get pure cultures of the *B. bacteriovorus* isolates, plaques must be re-streaked on the fresh double layer agar plates for 2 or more cycles^11^. The longer incubation time is also inevitable during the *B. bacteriovorus* purification process.

Reduction on the prey bacterial optical density can be easily observed within 24 hours in the liquid conditions in the presence of *B. bdellovibrio* ^15^, while incubating for up to 3 days are necessary for the visible lytic plaques on the prey lawn agar^11, 15^. This discrepancy of the predation efficacy on the liquid or solid medium needs to be further clarified. In other words, whether there was a lag time in the predation on solid condition or the emerged plaques were being overlooked need to be verified.

Here we evaluated the advantages of GFP labeled prey *S. entertidis* in the predation process under different conditions. Not only provides reliable trends through the cultural fluorescence intensity reduction, The GFP labeled *S. entertidis* also enhances the background of prey lawn, making it easier to track the subtle predation in time on the double-layer agar plate (Scheme 1). We believe that this practical method would reduce the time spent on bacterial isolation, purification, and enumeration of the predator *B. bacteriovorus*.

## MATERIALS AND METHODS

### Strains and culturing

Bacterial strains are detailed in Table 1. *S. entertidis* ATCC 13076 were typically grown in lysogeny broth for 12 hours, the pellets were centrifuged at 5000 rpm for 10 min, washed and resuspended with HEPES buffer (25 mM HEPES free acid, pH7.6) supplemented with 2mM Calcium Chloride.

**Table 1.** Strains of bacteria used in this study.

| Strains | Source or reference |
| --- | --- |
| **Prey** |  |
| *S. enteritidis* | ATCC 13076 |
| Fluorescent *S. enteritidis* | GFP labeled ATCC 13076, in this study |
| **Predator** |  |
| *B. bacteriovorus* HD100 | ATCC 15357 |
| *B. bacteriovorus* Bd4 | Soli, in this study |
| *B. bacteriovorus* Bd5 | Sewage, in this study |
| *B. bacteriovorus* Bd10 | Estuarine, in this study |
| *B. bacteriovorus* Bd16 | Soli, in this study |
| *B. bacteriovorus* Bd19 | Soli, in this study |

278 ATCC, American Type Culture Collection.

For culturing GFP labeled ATCC 13076, 10 μg/mL erythromycin was added as the selective pressure. As previously descripted^10^, *B. bacteriovorus* strains were grown on stationary phase prey *S. entertidis* ATCC 13076 in HEPES buffer (25 mM HEPES free acid, pH7.6) supplemented with 2mM Calcium Chloride at 30 °C with shaking at 200 rpm, after 24 hours, supernatants were filtered through the 0.45 μm syringe filter (Millipore SLHVR33RB) for at least 2 times to harvest pure predator cells.

### Predation in the liquid media

Predation experiment was conducted as described^13^ with some modifications. The prey-predator cocultures were prepared by mixing the prey GFP labeled *S. entertidis* ATCC 13076 and predator *B. bacteriovorus* isolates with the final concentration of 2 X 10^9^ CFU/mL and 2 X 10^7^ PFU/mL accordingly into 10 mL HEPES buffer (25 mM HEPES free acid, pH7.6) supplemented with 2mM Calcium Chloride. The cultures were shacking with 200 rpm at 30 °C, fluorescence intensity (Ex/m =395/509 nm) and the optical density at 600 nm were measured using Infinite M200 Microplate (Tecan) at 0, 2,4,8,12,24, and 36 hours.

### Growing plaques on the agar plate

Agar infusions containing 25 mM HEPES free acid, pH7.6, 2mM Calcium Chloride, and 0.6% agarose were kept at 55 °C, then either ATCC 13076 or GFP labeled ATCC-13076 was added into the media to a final concentration of 2 X 10^9^ CFU/mL. Then gently mixing and pouring it onto the plate. Different *B. bacteriovorus* isolates were inoculated in the congealed agar plate. The plates were incubated upside-down at the constant temperature and humidity chambers with 30 °C and 80% humidity conditions. Plaques on the background of different prey lawn were imaged under visible light and UV light (365 nm) using ChemiDox XRS+ Molecular Imager System (Bio-rad).

## RESULTS

### Fluorescence reflects the prey density during the predation process

In the predation experiment, suspensions containing *B. bacteriovorus* HD100 and GFP labeled *S. entertidis* ATCC 13076 was mixed under the initial predator/prey ratio of 1:100 at time zero, the fluorescence intensity (Ex/Em 395/509 nm) and the optical density at 600 nm were monitored regularly. As shown in Figure 1A and B, the optical density of prey suspension at 600 nm was decreased from 0.44 to 0.26 after predated for 36 hours, while the predator deleted group exhibited little reduction at the same time. By comparison, the fluorescence intensity of the HD100 treated group showed nearly 20-fold reduction from 19329 a.u. to 1154 a.u., which still exhibited a consistent decreasing trend with the optical density of prey suspension. A strong correlation (p<0.0001) between the fluorescent intensity and absorbance of the prey suspension was also suggested in Figure 1C. These results demonstrated that the reduction in optical density of prey suspension could be reflected as a significant fluctuation in the bacterial fluorescence intensity, thus indicating the presence and predation of the obligate predator *B. bacteriovorus* in real time.

**Figure 1.**
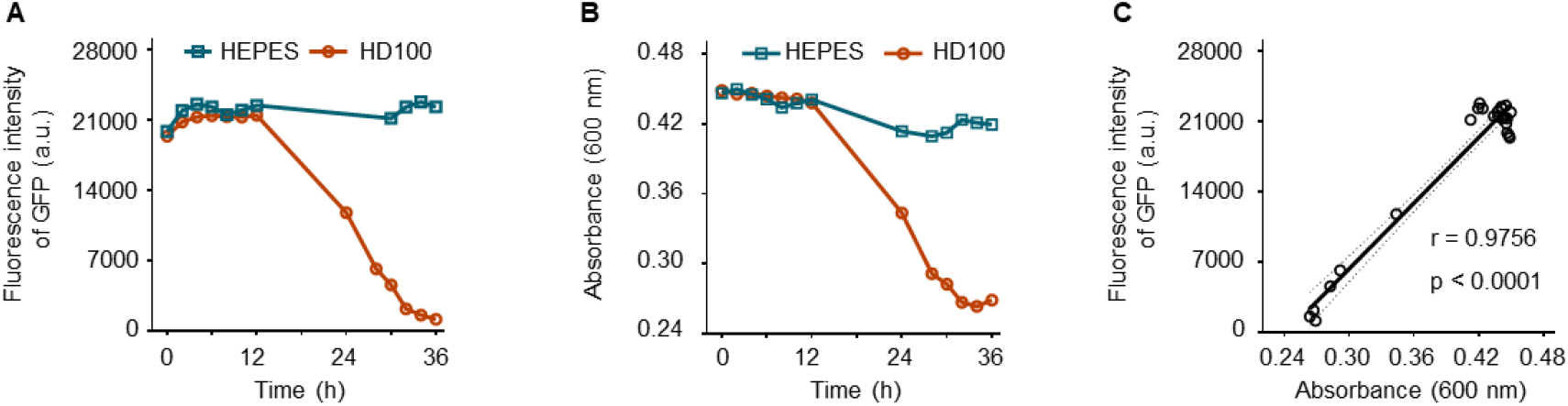
Fluorescence intensity and absorbance of the prey medium suggests the predation of *B. bacteriovorus*. A and B, the fluorescence intensity (A), and the absorbance (B) changing curves of GFP labeled pray *S. entertidis* ATCC 13076 in the presence of the predator *B. bacteriovorus* HD100 (red line) or HEPES (blue line). C, correlation analysis of fluorescent intensity and absorbance of bacterial medium.

### Fluorescence reduction reflects the lytic of prey cells

In this part, five clinical isolated *B. bacteriovorus* strains including Bd4, Bd5, Bd10, Bd16, and Bd19 were also used to verify the sensitivity of fluorescence on suggesting the bacterial density of prey populations. Although some slight variations, these *Bdellovibrio* isolates displayed the same predation trend with the type-strain HD100 (Figure 2A, B). The wide range of fluorescence reduction was larger than the absorbance changes during the predation assay. suggesting that data from the fluorescence was more reliable in the bacterial density measuring than the optical density. Besides, the predation by *B. bacteriovorus* isolates on *S. entertidis* reduced the viable prey population from the 10^9^ CFU/mL in the case of incubation without predator to less than 10^7^ CFU/mL after 36 hours in the case of incubation with different predator (Figure 2C, D), demonstrating that the reduction in prey fluorescence coincided with the lytic of the prey isolates. Though more than 99% viable prey was lysate by *B. bacteriovorus* after 36 hours in the liquid medium, 2–5 days incubation was always performed for allowing clear observation of plaques on the double-layer plate^11^, which would take longer time in the clinical isolation and purification of *Bdellovibrio*. Whether there was a lag time in the predation on solid condition or the emerged plaques were being overlooked need to be verified.

**Figure 2.**
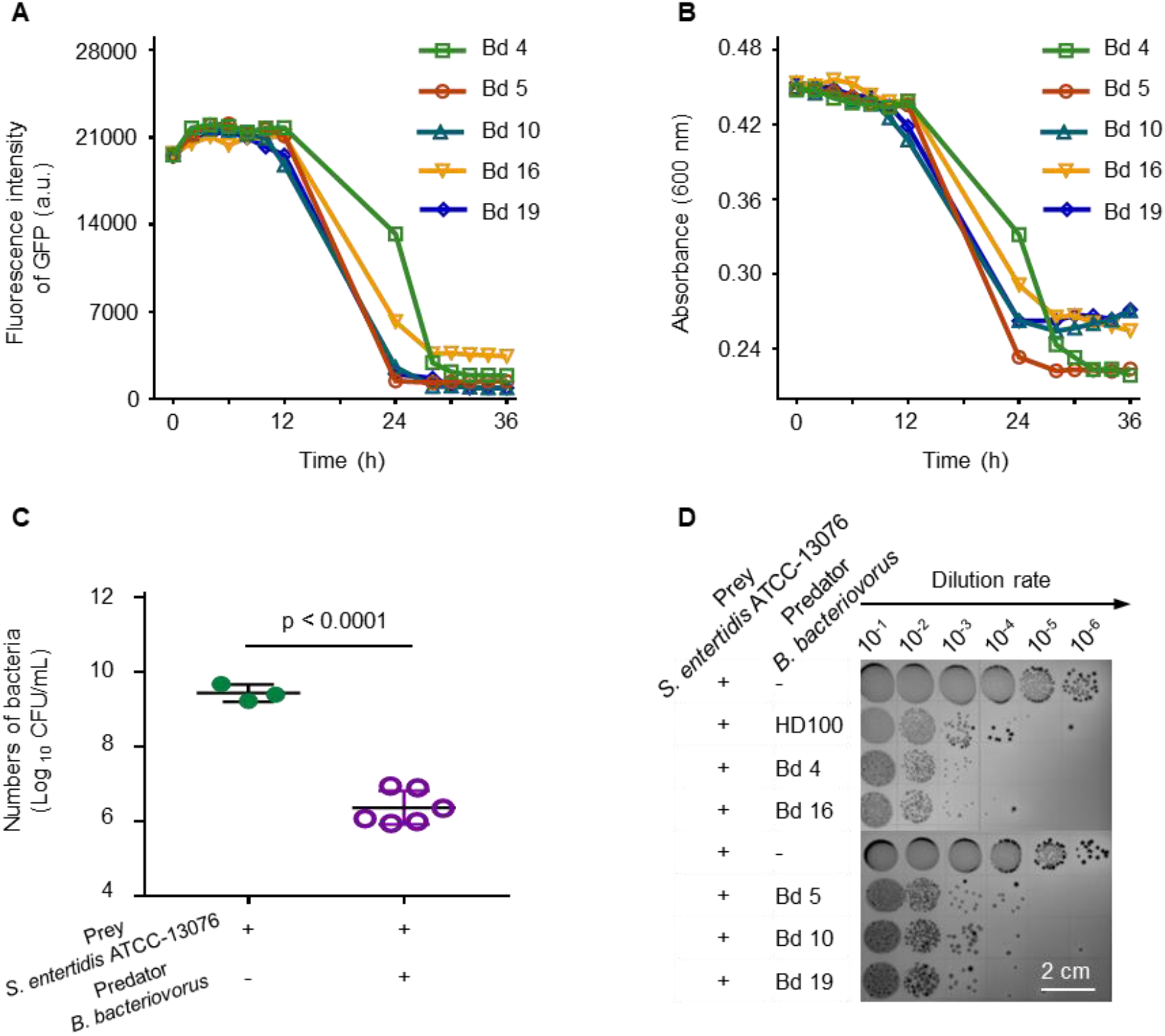
Viability of prey was significantly reduced with the predation of *B. bacteriovorus*. A and B, the fluorescence intensity and absorbance changing curves of pray medium in the presence of different predator isolates. C, predation efficiency of B. bacteriovorus strains on *S. entertidis*. D, representative images of viable *S. entertidis* ATCC 13076 over 36 h in HEPES buffer with or without *B. bacteriovorus* isolates.

### Plaques are visible under fluorescence-labeled prey lawn agar in the early stage

After streaking the obligate predator *B. bacteriovorus* HD100 on the double layer agar plate and incubating at 30 °C for 36 hours, no plaques were observed in either *S. entertidis* ATCC 13076 or GFP labeled ATCC 13076 contained agar under visible light (Figure 3A). However, when using the UV light, the *S. entertidis* ATCC-13076 contained lawn showed no fluorescence signal (Figure 3B), while the GFP labeled *S. entertidis* ATCC 13076 contained prey agar displayed an enhanced background due to the green light emission of GFP under ultraviolet light, and the regular plaques clearly illustrated the lysis of prey (Figure 3B). As shown in the inset image of Figure 3B, plaques exhibited significant reduction in gray value compared to the normal area. In other words, the plaques were already generated in the prey agar with the predation of *B. bacteriovorus* isolates for 36 hours, while these lytic plaques could not be found under visible light (Figure 3A). Nevertheless, when using the GFP-labeled prey, the predator existed area was easily observed as the regular lytic black plaques under the enhanced fluorescence background (Figure 3B). These individual plaques are clearly ready for enumeration or the next purification process of *B. bacteriovorus*.

**Figure 3.**
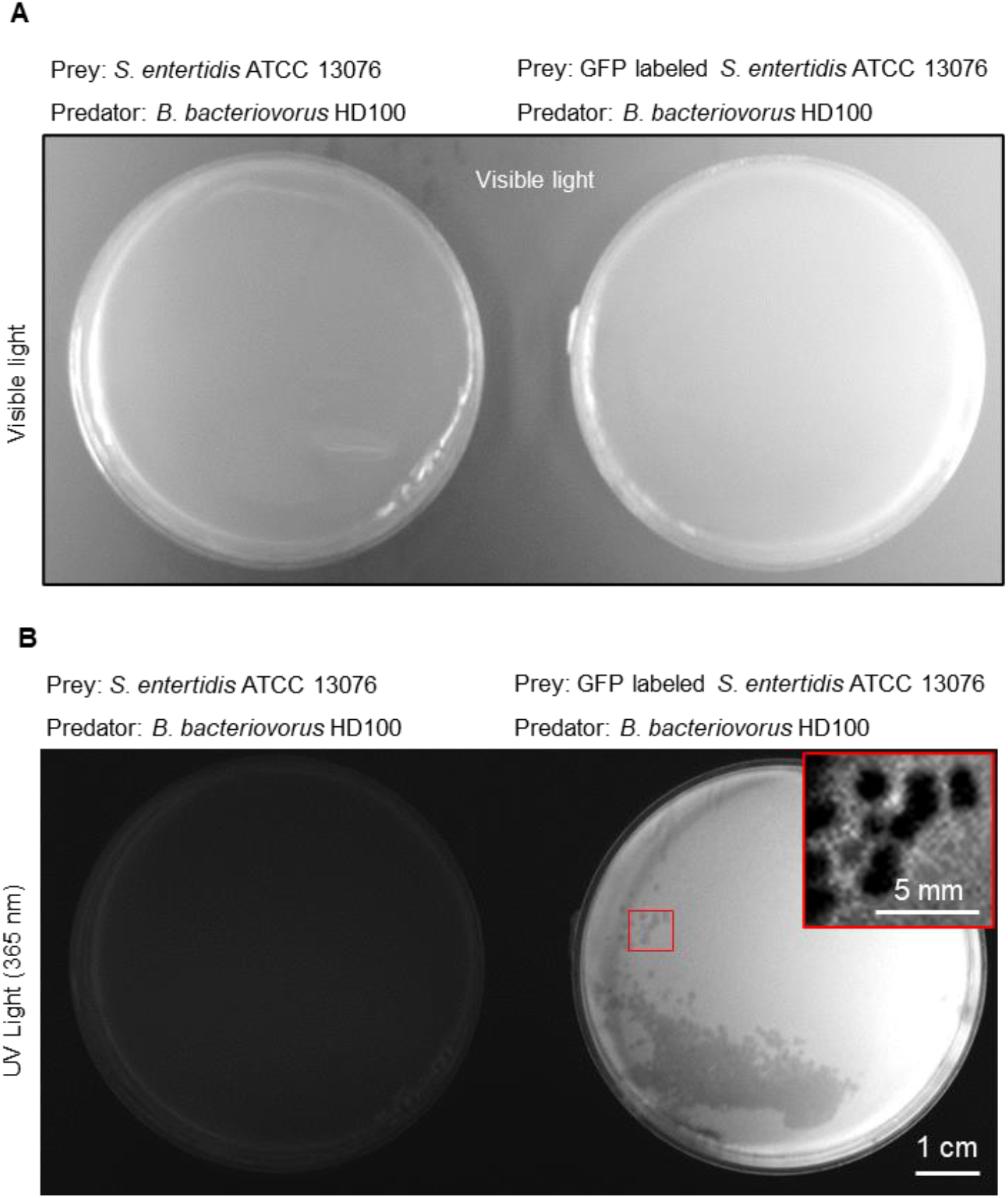
Plaques are visible under UV light (365 nm) on the double layer agar in the presence of fluorescent prey. A, *B. bacteriovorus* HD100 was scattered on the double-layer plate, the prey are *S. entertidis* ATCC 13076 (left) or GFP labeled *S. entertidis* ATCC 13076 (right) respectively, images were obtained under visible light. B, the same plate are imaged under UV light (365 nm).The insert image enlarged the plaques area of the red box.

## DISCUSSION

Since the time that the obligate predator *B. bdellovibrio* was isolated in 1960s, prey cells are always supplemented for the growth of the predatory bacteria. Though PFU enumeration on the double-layer agar are commonly applied for detect and quantify the predator populations^11^, this paradigm is a time-causing work. 2-5 days incubation are always performed during the isolation and purification processes of *Bdellovibrio spp*. Throughout the years, bioluminescent or fluorescent prey was applied to monitor the prey population during the predation process in time at liquid medium^7, 9^, which enormously move the field forward^17^. Advantages of the bioengineered prey in suggesting the predatory bacteria under solid condition also need to be clarified.

In our study, GFP was expressed in the prey strain *S. entertidis*. The reporter fluorescence enhanced the background of prey lawn in the agar plate, which could better track the subtle predation of predator *B. bacteriovorus*. Here, we confirmed that the plaques were already generated before we monitored it under visible light. Except for being utilized to monitor the predation and to investigate the specific role of defined prey genes in the predation process^18, 19^, the bioengineered prey can also be applied to visualize the predator in the earlier edge.

This work is not presented without flaws, though different *B. bacteriovorus* strains containing soil, water, and fecal sources were confirmed to be effective in predating *S. entertidis* ATCC 13076, more credible data need to be presented for validating the prey susceptibility to all *Bdellovibrio*. Besides, a precise timepoint that plaques appeared in the prey lawn need to be further explored, which might be helpful in further shorten the incubation time in the plate conditions.

## CONCLUSION

In the present study, a convenient, efficient method was introduced for promptly visualizing the plaques of *B. bacteriovorus* on the double layer agar plate. We confirmed that the plaques were already generated before we can observe it by eyes. When introducing GFP in the prey *S. e*ntertidis, the enhanced background of prey lawn allows us to track the subtle predation in time on the solid medium. This practical method reduces the time spent on bacterial isolation, purification, and enumeration of the predator *B. bacteriovorus*.

## ACKNOWLEDGEMENTS

This work was supported by the National Key Research and Development Program of China (2022YFD1801600) and National Nature Science Foundation of China (32230106)

## AUTHOR CONTRIBUTIONS

K.Z. designed and supervised this project. Q.Z. and J.X. performed the experiments. Q.Z., J.X., and K.Z. did the data analysis. Q.Z. and K.Z. wrote the manuscript.

## COMPETING INTERESTS

The authors declare no competing interests.

**Scheme 1.**
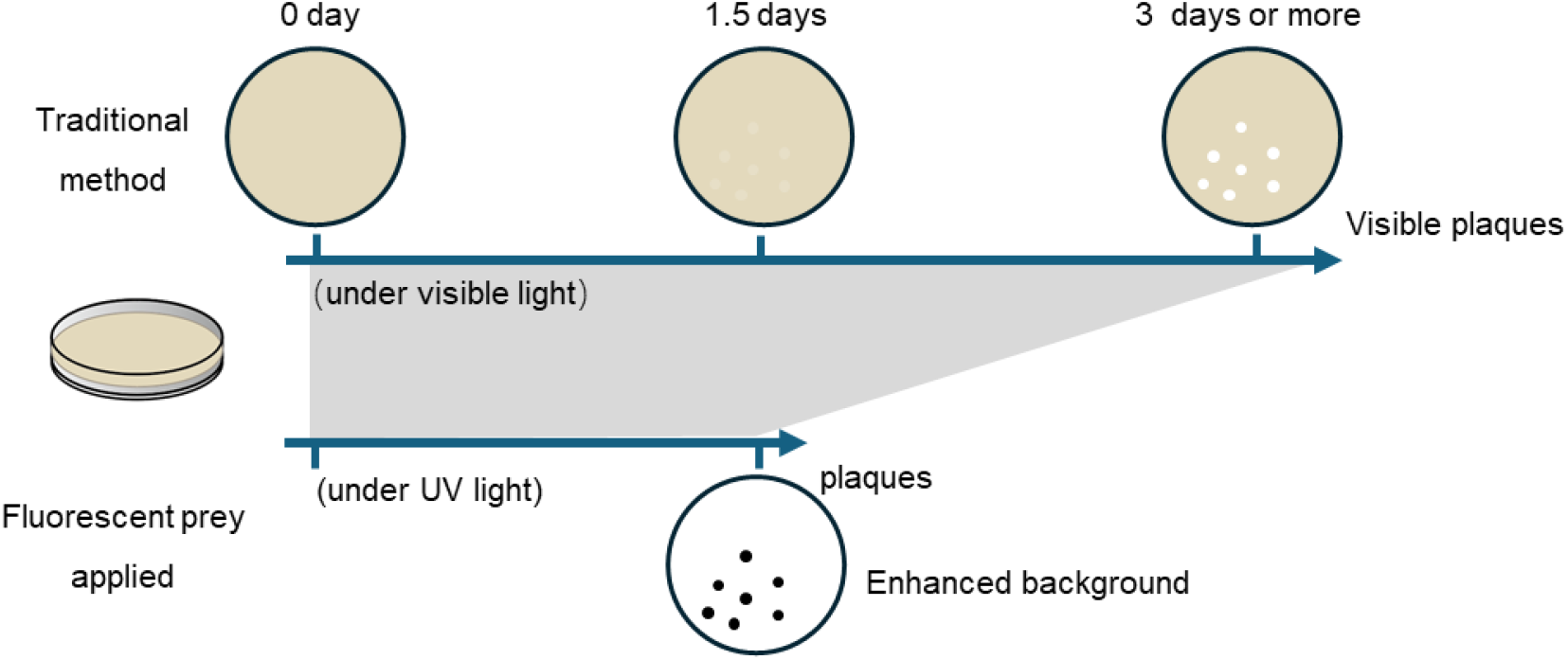
Schematic illustration of the fluorescent prey for plaques visualization. For growing plaques of *B. bacteriovorus*, 3 days are always necessary for observing lytic regular plaques, this time-consuming step reduces the efficiency of *B. bacteriovorus* purification processes. When using fluorescent prey, plaques are easily being observed on the enhanced background of prey lawn under UV (365 nm) light at day 1.5.

